# Factors that influence the thymic selection of CD8αα intraepithelial lymphocytes

**DOI:** 10.1101/761601

**Authors:** Nadia S. Kurd, Ashley Hoover, Jaewon Yoon, Brian M. Weist, Ellen A. Robey

## Abstract

Thymocytes bearing αβ T cell receptors (TCRαβ) with high affinity for self-peptide-MHC complexes undergo negative selection or are diverted to alternate T cell lineages, a process termed agonist selection. Among thymocytes bearing TCRs restricted to MHC class I, agonist selection can lead to the development of precursors that can home to the gut and give rise to CD8αα-expressing intraepithelial lymphocytes (CD8αα IELs). The factors that influence the choice between negative selection versus CD8αα IEL development remain largely unknown. Using a synchronized thymic tissue slice model that supports both negative selection and CD8αα̣IEL development, we show that the affinity threshold for CD8αα IEL development is higher than for negative selection. We also investigate the impact of peptide presenting cells and cytokines, and the migration patterns associated with these alternative cell fates. Our data highlight the roles of TCR affinity and the thymic microenvironments on T cell fate.

## INTRODUCTION

During development, thymocytes are tested for the ability of their T cell receptor (TCR) to recognize self-peptide bound to Major Histocompatibility Complex (MHC) proteins. While low to moderate self-reactivity is required for maturation and CD4/CD8 lineage commitment (positive selection), cells with high self-reactivity must be either deleted (negative selection) or diverted into alternate T cell lineages in order to maintain self-tolerance. Lineage diversion of self-reactive thymocytes (termed agonist selection)^1^ has been extensively studied for thymic-derived T_regs_ (tT_regs_) specific for MHC class II, and invariant natural killer T cells (iNK T cells) specific for the non-classical MHC I molecule CD1d. More recently, evidence has emerged that agonist selection by a variety of different MHC molecules, including classical and nonclassical MHC I, can lead to the development of thymic precursors to intraepithelial lymphocytes (IEL) that express TCRαβ and CD8αα (hereinafter referred to as CD8αα IELs)^2–5^. Although the exact roles of CD8αα̣IELs remain unclear, there is evidence that they, together with other IEL populations including TCRαβ+CD8αβ IEL derived from conventional CD8 T cells and TCRγδ+ IEL, help to maintain immune homeostasis and tolerance in the gut^6–8^.

While both negative selection and agonist selection are directed by a strong TCR signal, the factors that specify these divergent fates remain unclear. One popular notion posits that the degree of TCR self-reactivity required for agonist selection is intermediate between positive and negative selection. However, current evidence suggests that repertoire of tT_regs_ is quite broad, and overlaps extensively with both conventional CD4 T cells as well as with thymocytes undergoing negative selection^9, 10^. Early evidence that strong TCR signals in the thymus drive CD8αα IEL development came from mice expressing a defined MHC class I-specific TCR (OT-I) along with transgenic expression of the agonist ligand by medullary thymic epithelial cells (RIPmOVA mice)^5^. Further evidence came from the observation that the absence of the costimulatory CD28-B7 axis inhibited the negative selection of superantigen-reactive thymocytes, and favored their diversion into the CD8αα IEL lineage^11^. Recent studies using mice with monoclonal expression of TCRs cloned from endogenous CD8αα̣IELs provided evidence that CD8αα̣IEL precursors are found within a highly self-reactive thymocyte population enriched in thymocytes undergoing negative selection^2, 12^. Whether stronger TCR signals are required for CD8αα IEL development compared to negative selection remains unknown.

Another factor that can affect thymocyte fate is access to cytokines. For example, the related cytokines IL-2 and IL-15 play critical roles in the agonist selection of tT_regs_ and iNK T cells^13–16^. CD8αα IEL precursors express CD122, a shared subunit of the IL-15 and IL-2 receptors, suggesting the ability to respond to these cytokines^2, 4, 12, 17^. Although IL-15 is important for the survival and maintenance of mature CD8αα IELs in the gut, its role in their thymic development is less clear^8, 18–20^. One challenge in addressing this question is the lack of unambiguous markers for thymic CD8αα IEL precursors. For example, thymic IELp are often identified as TCR signaled TCRαβ+ CD4-CD8-thymocytes, however this may exclude IEL precursors with intermediate levels of CD4 and CD8^2, 12, 17^. In addition, NK1.1 is sometimes used to exclude iNKT cells from the “IELp” population, however this would also exclude subsets of IELp that express this marker^4, 17^. In addition, even after excluding iNKT cells and CD4 or CD8 expressing thymocytes, thymocytes defined as IEL precursors still exhibit considerable heterogeneity^4, 17, 21^, and it is unclear what fraction of these cells are bone fide precursors of mature CD8αα IELs.

Another unresolved question is the pattern of thymocyte motility and signaling during agonist selection. Previous studies have shown that negative selection is associated with migratory arrest and sustained TCR signals, whereas positive selection is associated with continued migration interspersed with pauses and transient TCR signals^22^. We previously examined the motility of surviving thymocytes in a steady state model of tolerance to a class I MHC-specific tissue restricted self-antigen (OT-I TCR and RIPmOVA transgenic mice), and reported a confined migration pattern of autoreactive thymocytes within zones of ∼30 microns diameter in the thymic medulla^23^. While that study did not address the ultimate fate of the autoreactive thymocytes, it is possible that the surviving thymocytes were undergoing agonist selection within confinement zones. However, the TCR signaling and motility patterns associated with agonist selection have not yet been specifically addressed.

In the current study we investigated these questions by examining a synchronous wave of CD8αα̣IEL development in thymic tissue slice cultures. We and others have previously used this experimental system to examine positive selection, negative selection, tT_reg_ development, and thymocyte migration^22–30^. We showed that thymocytes bearing a class I specific TCR (OT-I) undergo either negative selection or CD8αα̣IEL development when cultured on thymic slices in the presence of agonist ligand, and we used this system to investigate factors that influence whether autoreactive thymocytes undergo death or agonist selection. We found that higher affinity TCR signals were required for the development of CD8αα IELs compared to those required for negative selection. We also showed that IL-15 has an inhibitory effect on negative selection, and that both IL-2 and IL-15 promote the maturation and expansion of CD8αα IEL precursors. Agonist peptide presentation by both hematopoietic and non-hematopoietic cells could support CD8αα IEL development in this system, although hematopoietic cells were required to drive later stages of maturation, marked by T-bet induction and proliferation. We also observed that agonist selection was accompanied by elevated intracellular calcium and a confined migration pattern. Our results define key factors that determine whether self-reactive T cells undergo negative selection or adopt the CD8αα̣IEL fate in the thymus.

## RESULTS

### A proportion of self-reactive thymocytes do not undergo negative selection

To study factors that influence thymocyte fate following antigen encounter, we employed a thymic tissue slice system in which thymocytes of defined specificity are overlaid onto vibratome-cut thymic slices, after which they migrate into the slice and undergo a relatively synchronous wave of development^22, 25, 26^. In this study we used thymocytes from OT-I TCR transgenic mice and exposed them to agonist peptide derived from chicken ovalbumin (SIINFEKL, OVAp) by adding OVAp directly to tissue slices from wild type (WT) mice, or by using thymic slices from RIPmOVA transgenic mice, in which ovalbumin is expressed by a subset of medullary thymic epithelial cells. We evaluated the efficiency of negative selection in this system by including a reference population bearing an irrelevant TCR (F5 TCR transgenic or wild type thymocytes), and quantified thymocyte death as the ratio of live OT-I:reference thymocytes remaining in the slice. Consistent with previous studies^25, 31^, we observed a significant loss of OT-I thymocytes on both RIPmOVA slices and slices to which OVAp had been added, as well as a large proportion of viable OT-I thymocytes that remained in the slice after 16 hours of culture (Figure 1a). The majority of surviving OT-I thymocytes expressed high levels of CD69, indicating that the surviving thymocytes had encountered antigen and experienced strong TCR signals (Figure 1b).

**Figure 1.**
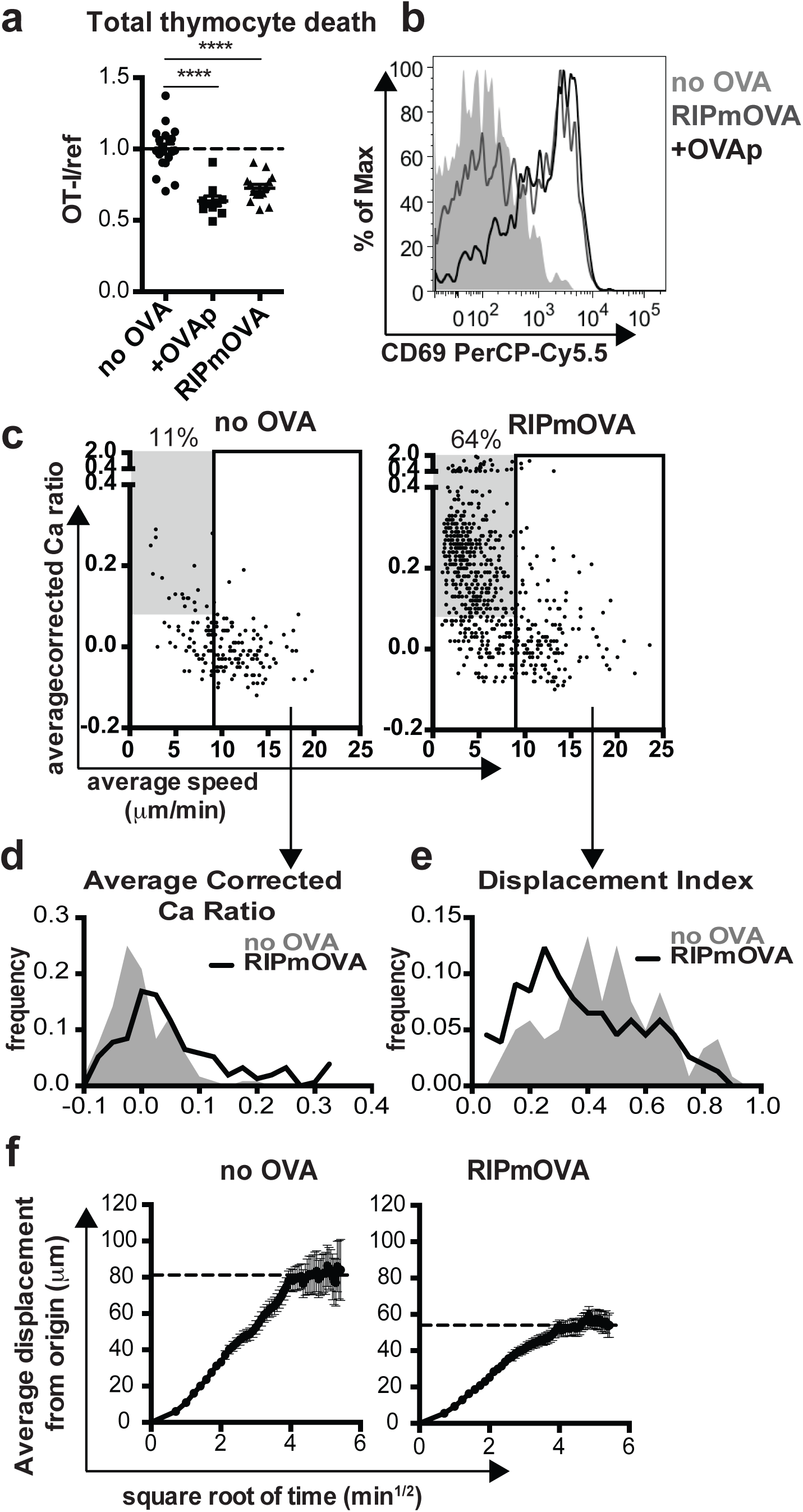
A proportion of OT-I thymocytes survive negative selection, and exhibit confined migration. (a,b) Total OT-I thymocytes were overlaid onto the indicated thymic slices with reference thymocytes, and slices were harvested after 16 hours for flow cytometric analysis. (a) Negative selection displayed as the ratio of live OT-I thymocytes relative to live reference thymocytes, normalized to no OVA controls. (b) CD69 expression on gated OT-I thymocytes. Data are representative of (b) or pooled from (a) 4 independent experiments, with mean and SEM of n=19-20 thymic slices, where each dot represents an individual slice. ****p<0.0001 (one-way ANOVA with Bonferroni’s correction). (c-f) OT-I thymocytes were depleted of mature CD8 single positives (Supplementary Figure S1) and labeled with the fluorescent ratiometric calcium indicator dye Indo-1 LR prior to overlay on WT or RIPmOVA thymic slices. Thymocytes were allowed to migrate into slices for 2 hours, then imaged by two-photon microscopy 0-7 hours later. Data are pooled from 4 (RIPmOVA) or 2 (WT) imaging runs. (c) Average speed versus relative calcium (calculated as described in Materials and Methods) where each dot represents an individual thymocyte track (RIPmOVA, n=757; WT, n=188). Numbers indicate the % of tracked thymocytes with low speed and high calcium tracks (grey box: average speed<9 m/min and average corrected calcium>0.08). Solid rectangle represents cutoff for fast tracks (average speed>9 m/min) used for the plots shown in d-f (d-e) Average corrected calcium ratio (d) and displacement index (e) of fast (>9 microns/minute) tracks, where n=154 (RIPmOVA) or 120 (WT) tracks. (f) Average displacement versus the square root of time for fast (>9 microns/minute) tracks on WT or RIPmOVA thymic slices. Black dotted lines indicate estimated plateau of confined migration, and error bars indicate SEM.

Previous studies of the *in situ* behavior of autoreactive thymocytes have revealed migratory arrest and calcium flux following addition of agonist peptide^22, 25^. On the other hand, continued migration within confinement zones was reported in a steady state model of AIRE-dependent negative selection^23^. To further explore the behavior of auto-reactive thymocytes undergoing death or survival, we performed 2-photon time-lapse imaging of OT-I thymocytes within RIPmOVA thymic slices. We first depleted OT-I thymocytes of the most mature CD8 single positive cells (CD8SP) (Supplementary Figure S1). We then labeled thymocytes with a fluorescent ratiometric calcium indicator dye, allowed them to migrate into RIPmOVA thymic slices, and performed 2-photon time-lapse imaging in the medulla after 3-7 hours of culture. On average, OT-I thymocytes in RIPmOVA slices displayed higher intracellular calcium levels (average corrected calcium ratio of 0.2 vs. 0.0, Figure 1c) and lower speed (6.0 vs. 10 m/min, Figure 1C) compared to thymocytes on wild type (no OVA) slices. Interestingly, we observed two distinct behaviors amongst thymocytes on RIPmOVA slices: a population with low speed and high calcium levels (483/757, 64% of total tracks), and a population with more rapid migration (Figure 1c).

In contrast, the majority of tracks on slices without OVA exhibited rapid migration, with only 20/188 tracks (11%) exhibiting low speed and high calcium levels (Figure 1c). Notably, even the rapidly migrating (>9 m/min) thymocytes in RIPmOVA slices displayed slightly higher intracellular calcium compared to those in wild type (no OVA) slices (0.1 versus 0.0 average corrected calcium ratio, Figure 1C,D). Moreover, the rapidly migrating OT-I thymocytes in RIPmOVA slices also displayed a lower directionality index, and reduced displacement from origin of migration (Figure 1 e, f) indicative of a more confined pattern of migration. In addition, visual inspection of tracks of OT-I thymocytes in RIPmOVA slices revealed multiple examples in which two or more confined tracks occupied the same 3-dimensional space, in close proximity to DC and AIRE-expressing mTECs (Supplementary Movies 1 and 2). Thus, thymocytes can make two distinct responses to encounter with agonist ligands: either migratory arrest with high intracellular calcium, or rapid, confined migration with slightly elevated calcium.

### Thymocytes that survive negative selection are TCRαβ CD8αα̣intraepithelial lymphocyte precursors

We next considered the possibility that the surviving thymocytes might give rise to a non-conventional agonist selected T cell lineage, such as CD8αα IELs. To determine whether the thymocytes that survived negative selection represented CD8αα IEL precursors, we investigated the phenotype of OT-I thymocytes that survived after 96 hours of culture on thymic slices. We depleted OT-I thymocytes of the most mature CD8SP (Supplementary Figure S1), and overlaid the remainder onto WT thymic slices with OVAp, or RIPmOVA slices, harvested at 96 hours, and analyzed by flow cytometry. OT-I thymocytes that survived negative selection proliferated within the slice and expressed markers associated with a CD8αα IEL precursor phenotype, including CD122, PD-1, T-bet, and the gut homing marker □_4_ □_7_ integrin^2–4, 8, 17^ (Figure 2a). In addition, OT-I thymocytes that had been exposed to OVA downregulated CD8β̣ relative to CD8α, implying higher expression of the CD8αα form of the co-receptor (Figure 2a).

**Figure 2.**
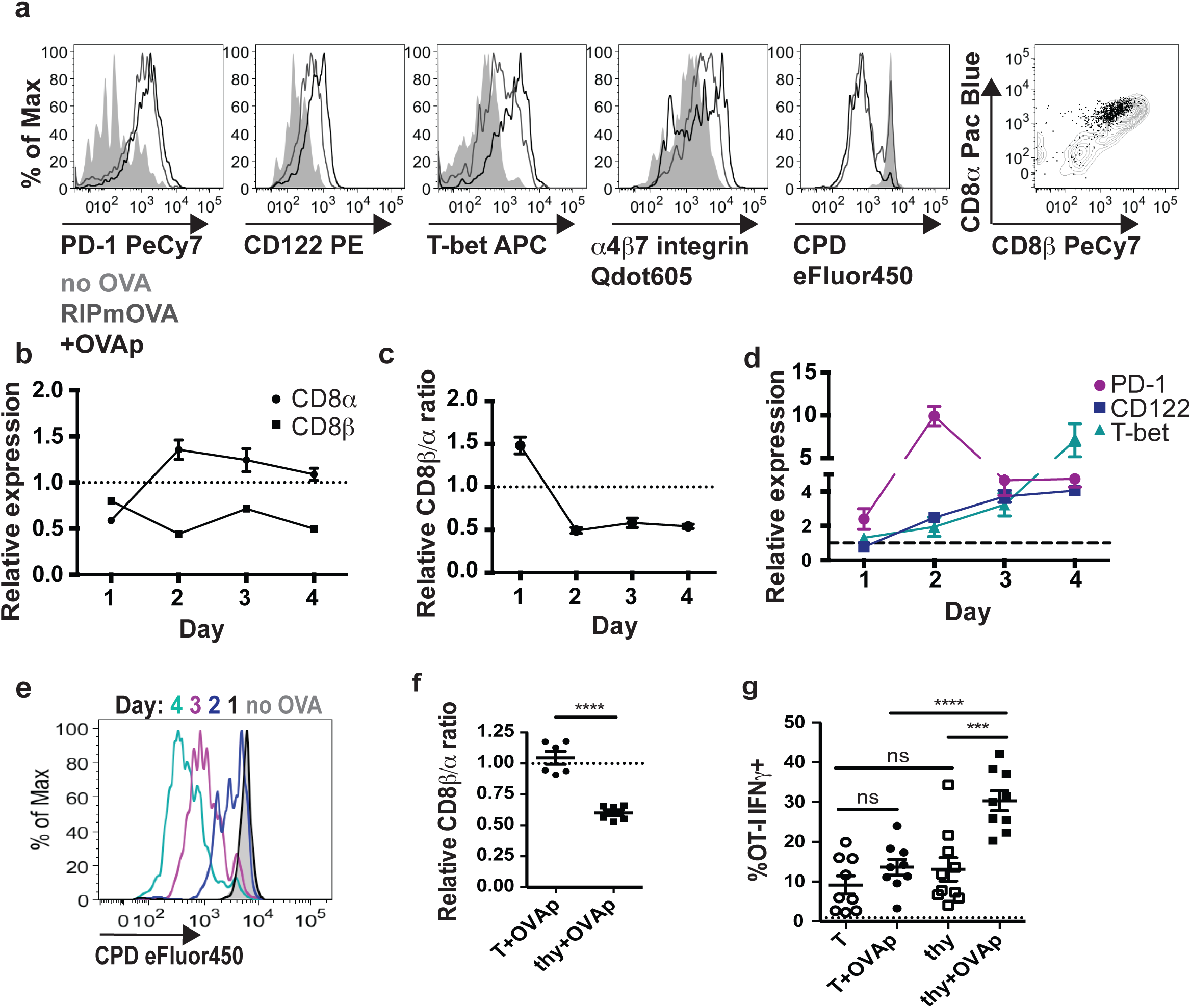
Thymocytes that survive negative selection develop a CD8αα IEL precursor phenotype. (a-e) OT-I thymocytes were depleted of the most mature CD8 single positives and stained with cell proliferation dye eFluor450 (CPD450) prior to overlay onto the indicated thymic slices, which were harvested for flow cytometric analysis 1-4 days later. Data are representative of 2 independent experiments. (a) Representative flow cytometry plots of CD122, T-bet, PD-1, □4□7 integrin, CPD450, and CD8α versus CD8βexpression in gated OT-I thymocytes cultured on RIPmOVA thymic slices, or WT thymic slices with or without OVAp for 4 days. Grey histograms and dots represent OT-I on slices without OVA, black lines and dots represent OT-I on slices with OVAp, and dark gray lines represent OT-I on RIPmOVA slices. (b) Expression of CD8α and CD8βby OT-I thymocytes cultured on WT thymic slices with OVAp for the indicated times, displayed as the ratio of the Mean Fluorescence Intensity (MFI) of sample relative to no OVA control slices. (c)Relative CD8β∼α̣ratio of OT-I thymocytes on WT thymic slices with OVAp over time, displayed as the ratio of CD8β MFI/CD8α MFI, normalized to no OVA control slices (d) Expression of T-bet, CD122, and PD-1 expression over time in thymocytes on WT slices with OVAp, displayed as the ratio of the MFI of the indicated marker relative to no OVA control slices. Data are compiled from two independent experiments, with mean and SEM shown. (e) Representative flow cytometry plot displaying dilution of CPD450 in OT-I thymocytes on slices with OVAp after the indicated days of culture. (f,g) OT-I thymocytes depleted of the most mature CD8 single positives or OT-I T cells collected from the lymph nodes were overlaid onto thymic slices to which OVAp was added, and analyzed 96 hours later (f) Fold change in CD8β∼α̣ratio of OT-I thymocytes (thy) or T cells (T) after 96 hours on thymic slices, displayed as the ratio of CD8β MFI/CD8α MFI relative to no OVA samples, where each dot represents an individual slice. Data are compiled from 2 independent experiments. (g) Production of IFNγby OT-I thymocytes (thy) or OT-I T cells (T) after restimulation with OVA-loaded DC, displayed as percent of IFNγ+ OT-I. Dashed line represents the average percent of IFNγ+ cells in culture with unloaded DC. Data are pooled from 3 independent experiments, with mean and SEM of n=9 thymic slices, where each dot represents an individual slice. ns not significant (p>0.05), ****p<0.0001 (one-way ANOVA with Bonferroni’s correction).

To determine the kinetics with which thymocytes undergo these phenotypic changes, we performed a time course in which thymic slices treated with OVAp were harvested after 1-4 days in culture. We observed that thymocytes exposed to OVAp exhibited decreased expression of both CD8β and CD8α̣at day 1 (Figure 2b), in line with studies demonstrating double-positive (DP) “dulling”, or the down-regulation of both CD4 and CD8, in response to antigen stimulation^2, 12, 32^. While CD8β expression remained low thereafter, expression of CD8α increased by day 2, resulting in a decreased CD8β to CD8α ratio (Figure 2c). We also observed that thymocytes exposed to OVAp exhibited a slight up-regulation of PD-1 as early as day 1, with maximal PD-1 expression at day 2 that declined, but remained elevated above baseline, thereafter (Figure 2d). In contrast, increases in CD122 and T-bet expression were not evident at day 1, but slight increases were apparent by day 2 that increased steadily thereafter (Figure 2d). Finally, we noted that the majority of proliferation occurred between 2-3 days (Figure 2e).

As an additional indication of the timing of phenotypic changes accompanying IEL development, we also examined the relationship between cell division and marker expression at day 2 of culture (Supplementary Figure S2a). Consistent with time course data, OT-I thymocytes upregulated PD-1 prior to cell division, whereas CD122 or T-bet upregulation occurred only on divided cells (Supplementary Figure S2b,c). Furthermore, CD122 and T-bet expression were detectable at division 1 and 2 respectively, suggesting that CD122 expression precedes T-bet expression (Supplementary Figure S2b,c). Additionally, OT-I thymocytes retained low expression of CD8β but increased expression of CD8α as they proliferated (Supplementary Figure S2d). These data indicate that a decrease in the CD8β̃α ratio and up-regulation of PD-1 begin very early after antigen encounter, followed by initiation of proliferation and increases in CD122, and finally T-bet expression.

To confirm that OT-I thymocytes that escaped negative selection were distinct from conventional activated T cells, we compared OT-I thymocytes to lymph node OT-I T cells after culture on thymic slices with or without OVAp. Both OT-I thymocytes and OT-I T cells proliferated extensively in OVA-containing slices, and had similar expression of CD122, PD-1, and T-bet (data not shown). However, whereas OT-I thymocytes exposed to antigen expressed lower levels of CD8β̣relative to CD8α, the relative expression of CD8β versus CD8α̣in OT-I T cells was unaffected by antigen exposure (Figure 2d). In addition, OT-I thymocytes that developed in the presence of OVA exhibited more robust production of IFNγ̣in response to restimulation by OVA-loaded DC (Figure 2g), a characteristic previously described for CD8αα̣IELs^33–35^. These observations indicate that OT-I thymocytes that survive negative selection are distinct from conventional CD8 T cells, and instead resemble CD8αα̣IELs.

To investigate whether OT-I thymocytes that survive negative selection represent bona fide CD8αα̣IEL precursors, we examined their ability to give rise to CD8αα IEL *in vivo*. We cultured allelically marked (CD45.1) OT-I thymocytes on thymic slices with or without OVAp for 96 hours, then dissociated the slices and transferred thymocytes into Rag2^-/-^ hosts (Figure 3a). One week after transfer, OT-I T cells, along with slice resident cells (CD45.2), could be detected in the intraepithelial lymphocyte compartment of the small intestine (Figure 3b). The percentage of OT-I thymocytes was reduced when donor populations came from cultures containing OVAp (Figure 3b), consistent with negative selection observed in these cultures (Fig 1a). Moreover, OT-I thymocytes that had been exposed to OVAp during thymic slice culture gave rise to a substantial population of CD8α+CD8β-IELs, whereas this population was minimal among cells that had not been exposed to OVAp (Figure 3c). Even amongst OT-I cells that retained some CD8β expression, the levels of CD8β relative to CD8α was lower amongst OT-I IELs that had not been exposed to OVA during development (Figure 3c). Taken together, these data demonstrate that OT-I thymocytes that survive negative selection in thymic tissue slices can home to the gut and give rise to CD8αα̣IEL.

**Figure 3.**
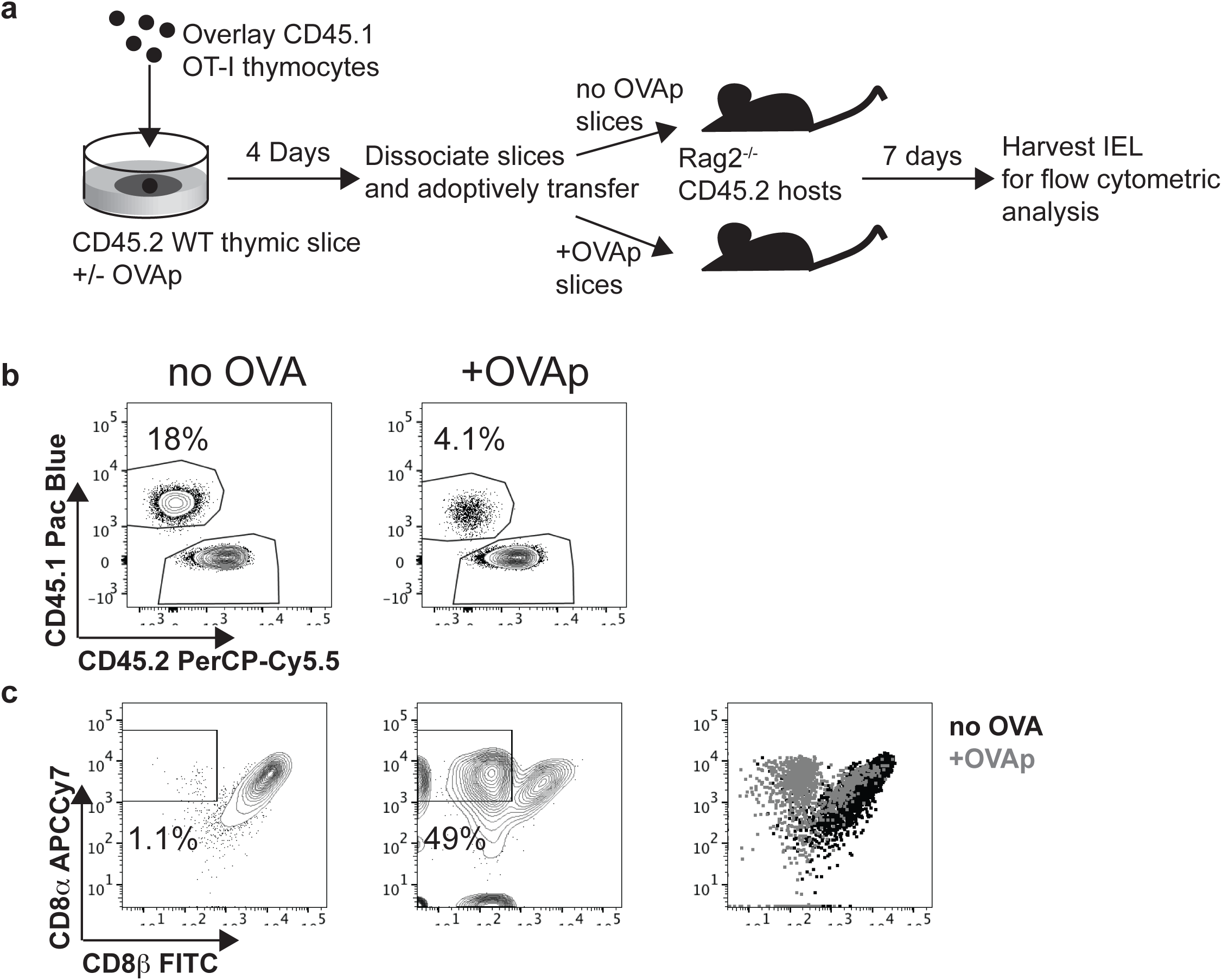
Thymocytes that survive negative selection can home to the intestinal epithelium and give rise to CD8αα IELs. (a) Schematic of experimental setup: OT-I thymocytes depleted of mature CD8 single positives were overlaid onto thymic slices that were treated with OVAp. After 96 hours, thymic slices were dissociated and adoptively transferred into Rag2^-/-^ mice. 1 week post-transfer, intraepithelial lymphocytes (IELs) were harvested from the small intestine. (b) Representative flow plots depicting gated TCR□+ IELs, showing the proportion derived from OT-I thymocytes versus slice resident thymocytes, based on CD45.1/CD45.2 congenic markers. (c) Representative flow plots depicting expression of CD8α versus CD8β in OT-I IELs. Gates show proportion of OT-I thymocytes that are CD8β-. Data are representative of 2 independent experiments, with n=2-4 mice per condition in each experiment.

### Higher affinity threshold for CD8αα̣IEL development compared to negative selection

Since both negative selection and agonist selection into the CD8αα̣IEL lineage occur within the thymic slice, this provides a useful experimental system for investigating factors that influence these alternative fates. Negative selection and CD8αα̣IEL development both require strong TCR signals compared to positive selection, but whether the TCR affinity threshold for CD8αα̣IEL development is higher or lower than that for negative selection remained unclear. To investigate this question, we added high affinity, SIINFEKL OVA peptide (OVAp), or a peptide variant with lower affinity for the OT-I TCR (Q4p)^36^ to thymic slices and determined the extent of negative selection and agonist selection in response to each peptide. Q4p was able to induce negative selection and stimulated CD69 up-regulation on the majority of surviving thymocytes (Figure 4a,b). However, thymocytes exposed to Q4p failed to upregulate T-bet, CD122, and PD-1, and did not have the relative decrease in the ratio of CD8β̣to CD8α observed in response to OVAp (Figure 4c-f). Thus, although both OVAp and Q4p were able to induce negative selection, only the highest affinity OVA peptide was able to induce a CD8αα̣IEL precursor phenotype.

**Figure 4.**
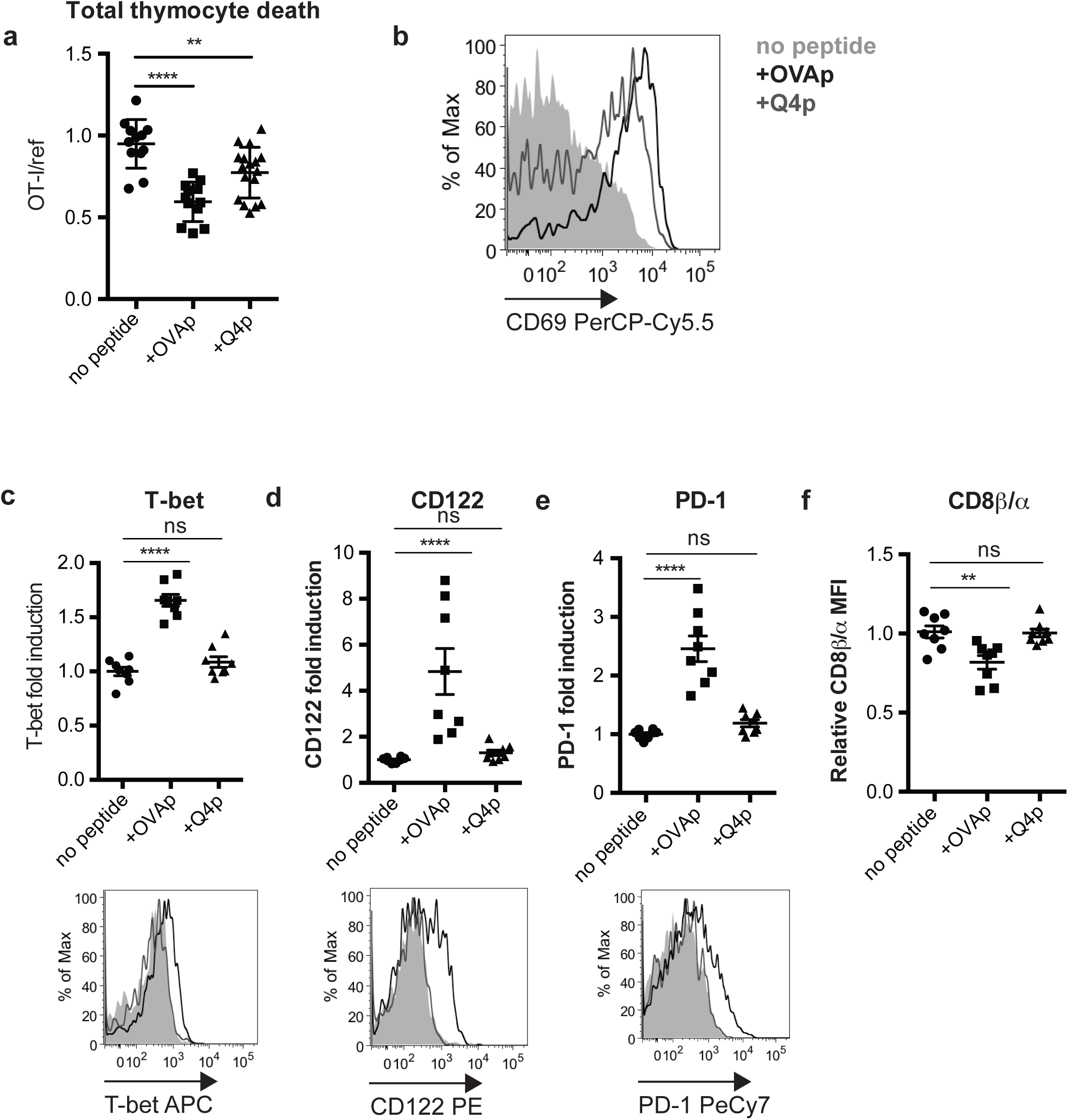
Only very high affinity agonists drive CD8αα IEL development. OT-I thymocytes depleted of mature CD8 single positives were overlaid onto WT thymic slices to which OVAp (SIINFEKL) or lower-affinity Q4 peptide was added. In some cases (a,b) a reference population was added with the OT-I thymocytes. Slices were harvested at 16 (a,b) or 96 (c-f) hours for flow cytometric analysis. (a) Negative selection displayed as the ratio of live OT-I thymocytes relative to live reference thymocytes, normalized to no OVA controls. (b) Antigen recognition of OT-I thymocytes displayed as expression of the activation marker CD69. Data are representative of (b) or pooled from (a) 3 independent experiments, with mean and SEM of n=11-15 total slices per condition, where each dot represents an individual slice. (c-e) Expression of T-bet (c), CD122 (d), or PD-1 (e) in OT-I thymocytes presented as fold induction displayed as MFI relative to levels in no peptide controls (top), or representative flow cytometry plots (bottom). (f) Fold change in CD8βαratio displayed as the ratio of CD8α MFI/CD8β MFI relative to no peptide samples, where each dot represents an individual slice. Data are representative of (bottom panels) or pooled from (top panels) 2 independent experiments, with mean and SEM of n=6 total slices per condition, where each dot represents an individual slice. ns not significant (p>0.05), **p<0.01, ****p<0.0001 (one-way ANOVA with Bonferroni’s correction).

### IL-15 prevents negative selection while both IL-15 and IL-2 promote CD8αα IELp maturation

In addition to CD122, a shared subunit of the IL-15 and IL-2 receptors, OT-I thymocytes exposed to OVAp also up-regulated CD25, the high-affinity □-chain of the IL-2 receptor (Supplementary Figure S3) suggesting that both cytokines might influence thymocyte fate. To examine this question, we added exogenous IL-15 or IL-2 to thymic slices containing OT-I thymocytes and OVAp, and investigated the effect on negative selection and CD8□□ IEL development. To supply IL-15, we used IL-15/IL-15Rα complexes (IL-15c) to mimic trans-presentation of IL-15 bound to the IL-15Rα^37–39^. To supply IL-2, we used complexes of IL-2 and antibody specific for IL-2 (IL-2c), which have greater biological activity compared to IL-2 alone^40^. To assess effects on negative selection, we overlaid OT-I and reference thymocytes onto thymic slices followed by treatment with OVAp with and without added cytokines. We observed that IL-15, but not IL-2, decreased negative selection, as indicated by an increase in the OT-I:reference ratio at 16 hours (Figure 5a). This increase was not due to IL-15-induced proliferation, because OT-I thymocytes had not proliferated at this time point under any of the conditions (Figure 5a, data not shown). These data suggest that IL-15, but not IL-2, has an inhibitory effect on negative selection, and might promote the CD8αα IEL fate in part by limiting the death of self-reactive thymocytes that could otherwise develop into CD8αα IEL precursors.

**Figure 5.**
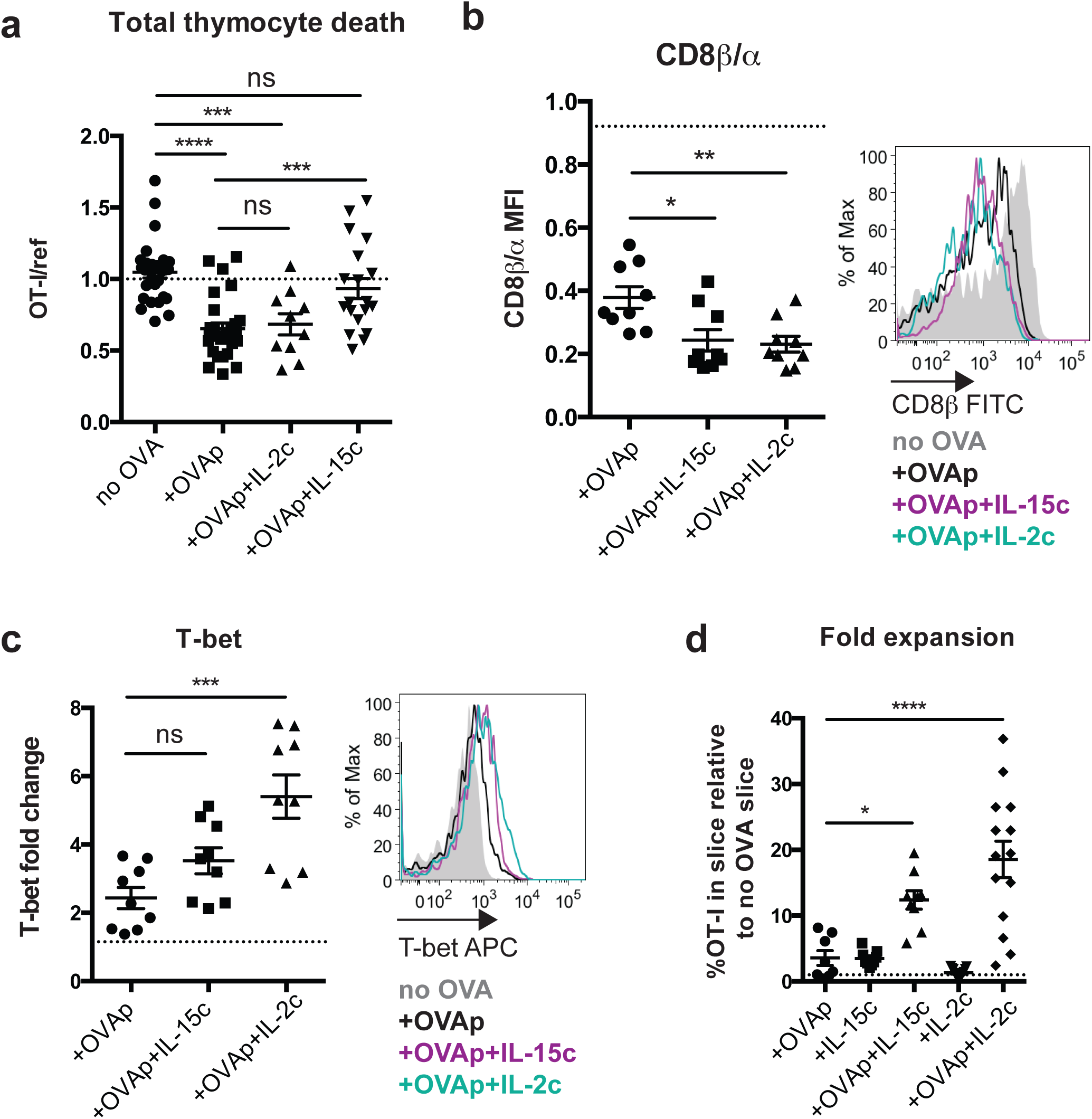
Distinct effects of IL-15 and IL-2 on survival, maturation, and expansion of CD8ααIEL precursors. Total OT-I thymocytes (a) or OT-I thymocytes depleted of mature CD8 single positives (b-d) were overlaid onto thymic slices to which OVAp was added, followed by treatment with IL-15Rα/IL-15 complexes or IL-2/α-IL-2 antibody complexes. Slices were harvested at 16 (a), 48 (b,c) or 96 (d) hours for flow cytometric analysis. (a) Negative selection displayed as the ratio of live OT-I thymocytes relative to live reference thymocytes, normalized to no OVA controls. Data are pooled from 3 independent experiments, with mean and SEM of n=13-19 total slices per condition, where each dot represents an individual slice. (b) Changes in CD8β̣expression displayed as the ratio of CD8β MFI/CD8α MFI relative to no OVA control samples (left) or representative flow cytometry depicting expression of CD8β(right). Dashed line indicates the average CD8β/CD8α ratio in no OVA control samples across all treatment conditions; no significant differences in the CD8β/CD8α ratio were observed in response to cytokine treatment without OVAp. (c) Expression of T-bet in OT-I thymocytes presented as representative flow cytometry (right), or fold induction of T-bet displayed as MFI relative to no OVA control samples (left). Dashed line indicates the average T-bet expression in no OVA control samples across all treatment conditions; no significant differences in T-bet expression were observed in response to cytokine treatment without OVAp. Data are representative of (right panels of b,c), or pooled from (left panels of b,c) 3 independent experiments, with mean and SEM of n=9 total slices per condition, where each dot represents an individual slice. (d) Fold expansion of OT-I thymocytes at 96 hours, displayed as percent of the total slice made up of OT-I thymocytes relative to untreated, no OVA control slices. Data are pooled from 4 independent experiments, with mean and SEM of 15-16 total slices per condition, where each dot represents an individual slice. ns not significant (p>0.05), *p<0.05, **p<0.01, ***p<0.001, ****p<0.0001 (one-way ANOVA with Bonferroni’s correction).

We also observed that treatment with both exogenous IL-15 and IL-2 resulted in a marked decrease in the ratio of CD8β̣to CD8α at 48 hours, as well as an increase in T-bet expression, which was more pronounced in response to IL-2 (Figure 5b,c). Additionally, both IL-15 and IL-2 resulted in a robust increase in the frequency of OT-I thymocytes present in the slice at 96 hours (Figure 5d). Taken together, these data suggest that both IL-15 and IL-2 signals can promote the CD8αα IEL phenotype and expand the CD8αα IEL precursor population.

### Peptide presentation by hematopoietic cells promotes the maturation and expansion of CD8αα IEL precursors

The thymus contains a variety of peptide-presenting cells, but which cells can serve as antigen presenting cells for CD8αα IEL development remained unclear. To address this question, we made use of Kbm1 mice, which carry an allelic version of the MHC class I K gene that is unable to stimulate OT-I T cells^41^. To restrict antigen presentation to hematopoietic cells (including DCs and macrophages), we transferred WT bone marrow into Kbm1 hosts (referred to as WT➔Kbm1 mice). Reciprocally, we transferred Kbm1 bone marrow into WT hosts to restrict antigen presentation to stromal cells (including epithelial cells) (referred to as Kbm1➔WT mice). We then compared the efficiency of CD8αα IEL development of OT-I thymocytes overlaid onto thymic slices from chimeric mice and treated with OVAp. We observed a modest reduction in the percent of OT-I thymocytes that expressed high levels of CD122 and PD-1 at 48 hours in both WT➔Kbm1 and Kbm1➔WT thymic slices, suggesting that both stromal and hematopoietic compartments contribute to the development of IEL precursors (Figure 6a). Interestingly, thymocytes that encountered antigen on hematopoietic cells proliferated more extensively (slices from WT➔Kbm1 and WT➔WT) than thymocytes that encountered antigen only on stromal cells (Kbm1➔WT slices) (Figure 6b). In contrast, upregulation of T-bet was significantly abrogated when hematopoietic cells could not present OVA (in Kbm1➔WT slices), while restricting antigen presentation to hematopoietic cells resulted in an increase in T-bet expression (in WT➔Kbm1 slices) (Figure 6c). Taken together, these data suggest that while peptide presentation by stromal cells can promote the initial differentiation of CD8αα IEL precursors, peptide presentation by hematopoietic cells more efficiently supports their late maturation, including proliferation and T-bet up-regulation.

**Figure 6.**
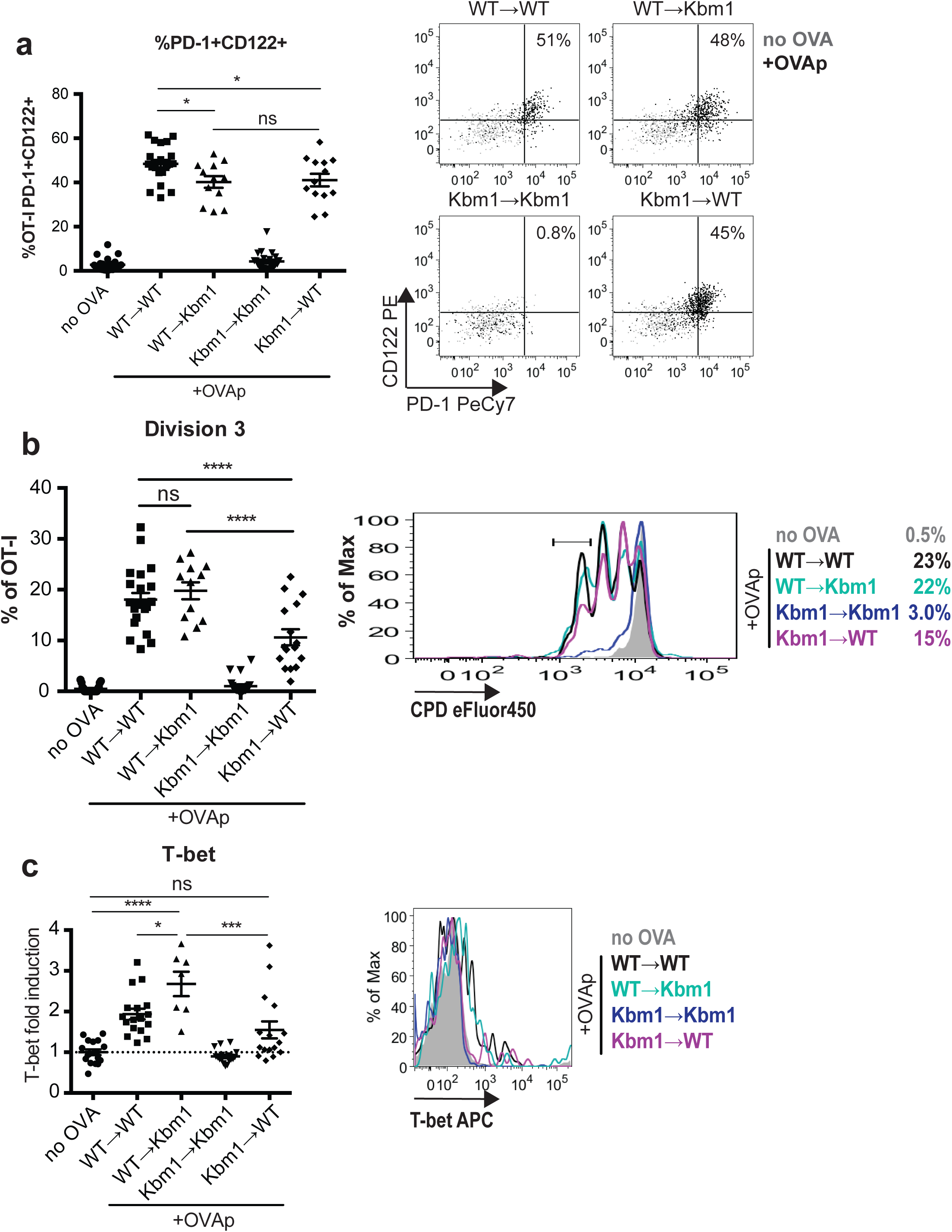
Effects of peptide-presenting cell on the development of CD8ααIEL precursors. OT-I thymocytes depleted of mature CD8 single positives were stained with CPD450 and overlaid onto thymic slices from WT-➔Kbm1, Kbm1➔WT, WT➔WT, or Kbm1➔Kbm1 bone marrow chimeric mice. In some experiments, WT or Kbm1 mice were used instead of WT➔WT or Kbm1➔Kbm1 control chimeras, respectively, with equivalent results. OVAp was added to thymic slices, which were harvested 48 hours later for flow cytometric analysis. (a) Percent of OT-I thymocytes expressing CD122 and PD-1, displayed as values (left) or representative flow cytometry plots (right). (b) Expression of T-bet in OT-I thymocytes exposed to OVAp presented as Mean Fluoresence Intensity (MFI) of T-bet relative to no OVA controls (left), or representative flow cytometry plot (right). Dashed line indicates the average T-bet expression in no OVA control samples across all treatment conditions (c) Proliferation of OT-I thymocytes displayed as percent of OT-I thymocytes that have undergone 3 cellular divisions, determined based on dilution of CPD450 according to the gate shown in the representative flow cytometry plot (right). Data are pooled from (left panels) or representative of (right panels) 3 (WT➔Kbm1 and Kbm1➔WT conditions) or 5 (no OVA, WT➔WT, Kbm1➔Kbm1 conditions) experiments with mean and SEM of n=12-22 total slices, where each dot represents an individual slice. ns not significant (p>0.05), *p<0.05, ***p<0.001, ****p<0.0001 (one-way ANOVA with Bonferroni’s correction).

## DISCUSSION

Strong TCR signaling during thymic development can lead to either the removal of self-reactive cells by negative selection, or their diversion into regulatory lineages, termed agonist selection. While this phenomenon has been extensively investigated for regulatory T cell development from MHC class II-restricted thymocytes, less is know about the agonist selection of CD8αα IEL. Here, we develop an experimental system in which thymocytes bearing an MHC class I-specific TCR undergo both negative selection and CD8αα IEL development after encounter with agonist peptide within thymic tissue slice culture, and we use this system to investigate factors that influence auto-reactive thymocyte fate. We found that higher affinity interactions are required to drive agonist selection of CD8αα IELs compared to that required for negative selection. We also found that both IL-2 and IL-15 promote the maturation and expansion of CD8αα IEL precursors, while IL-15 has the additional effect of inhibiting negative selection. Finally, our data suggest that while peptide presentation by both stromal and hematopoietic cells can drive the initial development of CD8αα IELs, hematopoietic cells more efficiently drives their expansion and further maturation. These findings identify factors that support CD8αα IEL development (Supplementary Figure S4), and highlight the importance of the thymic environment and peptide-presenting cell, along with TCR affinity for self-ligands, in shaping T cell fate.

By definition, agonist selection requires stronger TCR signals compared to positive selection, but the relative TCR signal strength required for agonist versus negative selection is less clear. For class II restricted T cells, recent studies indicate that a broad range of self-reactivity can lead to regulatory T cell development, resulting in a tT_reg_ TCR repertoire that overlaps extensively with both conventional CD4 T cells and thymocytes undergoing negative selection^9, 42^. In contrast, we find that CD8αα IEL development requires stronger TCR signals compared to negative selection, since a peptide with moderately high affinity for the TCR was capable of inducing negative selection, but not CD8αα̣IEL development. The notion that stronger TCR signals are required to drive CD8αα̣IEL development compared to negative selection is also consistent with the lack of overlap between the CD8αα IEL and conventional CD8 TCR repertoires^2, 12^ (Supplementary Figure S4a). The less stringent avidity requirements for development of tT_regs_ compared to CD8αα IEL may relate to the greater impact of cell death in thymic selection of class I versus class II restricted T cells^43^, and the relatively modest role of deletion in maintaining tolerance within CD4 lineage T cells due to the diversion of moderately self-reactive CD4 T cells into the regulatory T cell lineage^44, 45^. These findings might also help to explain the observation that class I-restricted T cells with moderate affinity for self are the primary mediators of autoimmunity^46^, since such thymocytes are more likely to escape negative selection, but less likely to undergo agonist selection, compared to thymocytes with higher affinity for self (Supplementary Figure S4c).

Taking advantage of the synchronicity of development in the thymic slice model, we define the kinetics of phenotypic changes in thymocytes undergoing CD8αα IEL development, revealing an initial transient up-regulation of PD-1 at 1 day and the induction of T-bet and proliferation at 3 days after agonist encounter. It is interesting to consider this phenotypic progression in light of current information about heterogeneity amongst thymic IEL precursors. A recent study identified PD1+T-bet- and PD-1-Tbet+ populations, and provided evidence that these correspond to distinct lineages, both of which can contribute to the mature IEL compartment ^4^. Other studies have proposed a precursor-product relationship between T-bet- and T-bet+ precursors, based on the more mature phenotype of T-bet+ IELp, as well evidence that in vitro exposure to IL-15 can induce T-bet upregulation in IELp^4, 8, 17, 21^. Importantly, these two possibilities are not mutually exclusive. Indeed, our data, together with previous reports, suggest that the PD-1+Tbet-compartment may be heterogeneous, with some cells being competent to leave the thymus and seed the intestinal compartment, and others destined to remain in the thymus and undergo further maturation into PD-1-T-bet+ IELp.

IL-15 plays a crucial role for CD8αα IEL in the intestine, but its role in the thymic development of CD8αα IEL precursors has not been fully resolved. IL-15^-/-^ mice have normal numbers of total CD8αα IEL precursors, but lack a subset of the most mature T-bet+ CD8αα IEL precursors ^4, 8, 21^. Here, we show that IL-15, as well as the related common gamma chain cytokine IL-2, can drive multiple aspects of CD8αα IEL development, including proliferation, CD8β down-regulation, and T-bet upregulation. In addition, IL-15, but not IL-2, can inhibit negative selection, suggesting that IL-15 signals might promote the CD8αα IEL fate both inhibiting negative selection and by actively promoting the CD8αα IEL developmental program (Supplementary Figure 4b). Interestingly, there is evidence that IL-2 both prevents negative selection of tT_regs_, and promotes their differentiation^47^. This could reflect a difference in the response of class I versus class II restricted thymocytes to IL-2, or a difference in the experimental systems used.

We have previously shown that distinct T cell fates are correlated with distinct patterns of thymocyte motility and TCR signaling^22, 24, 25^. For example, positive selection is characterized by transient TCR signals interspersed with periods of active migration, whereas negative selection is characterized by sustained TCR signals and migratory arrest. Here, we observed that thymocytes in a synchronous model of exposure to a tissue-restricted antigen exhibited two distinct patterns of motility, correlating with their two distinct fates in this system. Some thymocytes exhibited behavior consistent with negative selection (high intensity TCR signals and migratory arrest), whereas others displayed a distinctive motility behavior that might correlate with agonist selection. Specifically, these cells remained motile while exhibiting higher intracellular calcium levels and a more confined pattern of migration compared to thymocytes cultured without antigen. We also observed examples of “confinement zones”, in which the migration of multiple thymocytes appeared to be confined within the same 3-dimensional space. A similar pattern of confined migration was observed among thymocytes in the presence of a tissue-restricted antigen at steady-state ^23^. Thus, both the surviving thymocytes in a steady-state model of negative selection, and a portion of thymocytes in a synchronized negative selection, display a migration pattern suggestive of niches within the medulla that might support agonist selection.

Our observation that both peptide presentation by hematopoietic cells, as well as exogenous IL-2 and IL-15, enhanced certain aspects of the CD8αα IEL developmental program, suggest that thymic microenvironments enriched in these factors would support CD8αα IEL development. Medullary thymic epithelial cells (mTECs) are the major source of IL-15 in the thymus and closely associate with hematopoetic cells such as thymic DC^29, 48, 49^. Thymic IL-2 derives from hematopoetic cells, particularly thymocytes^50^, and there is also evidence that spatial linkage of IL-2 with antigen presenting cells such as DC promotes tT_reg_ development^10, 26^. Thus, high-affinity TCR ligand provided by DCs, together with trans-presentation of IL-15 by neighboring mTECs and IL-2 production by mature thymocytes receiving strong TCR signals, together might provide specialized niches for agonist selection in the medulla. Previous results also suggest that thymic microenvironments low for CD28 ligands would be expected to favor CD8αα IEL development over negative selection^11^. Other cytokines, such as TGFβ also contribute to CD8αα IEL development^51^, and could also contribute to an agonist selection niche.

Altogether, our data suggest that a “niche” that efficiently supports CD8αα̣IEL development includes presentation of high affinity self-peptides by hematopoietic-derived cells together with IL-15 and/or IL-2, and support the idea that confined migration within such niches is a characteristic of agonist selection. These findings help to elucidate mechanisms by which divergent thymocyte fates are achieved from the same initial stimulus, and should inform future studies aimed at understanding the requirements for CD8αα IEL development in the thymus.

## METHODS

### Mice

All mice were maintained and bred in-house in an American Association of Laboratory Animal Care-approved facility at the University of California, Berkeley. All procedures were approved by the University of California, Berkeley Animal Use and Care Committee. C57BL/6, C57BL/6-Tg(Ins2-TFRC/OVA)296Wehi/WehiJ (RIPmOVA), B6(Cg)-Rag2tm1.1Cgn/J (Rag2^-/-^), B6.C-H2-Kbm1/ByJ (Kbm1), and B6.SJL-Ptprca Pepcb/BoyJ (CD45.1) were from Jackson Labs. OT-I Rag2^-/-^ mice were from Taconic Farms. OT-I Rag2^-/-^ CD45.1 mice were generated by crossing OT-I Rag2^-/-^ mice to CD45.1 mice. F5 Rag1^-/-^ mice^52^, CD11cYFP mice^53^, and AireGFP mice^54^ have been previously described. CD11cYFP RIPmOVA mice and AireGFP RIPmOVA mice were generated by crossing CD11cYFP or AireGFP mice to RIPmOVA mice, respectively. To generate bone marrow chimeras, bone marrow was collected from the tibias and femurs of donor mice, then treated with ammonium chloride–potassium bicarbonate buffer to lyse red blood cells. Host mice were lethally irradiated (900rads) prior to intravenous injection of 5·10^6^ bone marrow cells.

### Thymocyte enrichment and labeling

Thymocytes were collected by dissociating whole thymic lobes through a 70 m cell strainer to obtain a cell suspension. For negative selection experiments, total OT-I and reference (B6 or F5 Rag1^-/-^) thymocytes were stained with Cell Proliferation Dyes eFluor450 (1□M) or eFluor670 (0.5□M) (Thermo Fisher Scientific) at 10^7^ cells/ml in PBS for 15 minutes at 37°C prior to overlay on thymic slices. For all other experiments, OT-I thymocytes were depleted of mature CD8 single positives using anti-human/mouse □7 integrin antibody (FIB504, Biolegend) and the EasySep Biotin Positive Selection Kit (Stemcell Technologies) according to product instructions. Following depletion, thymocytes were stained with CPD eFluor450 or eFluor670 as described above.

### Thymic slices

Preparation of thymic slices has been previously described in detail^55^. Briefly, thymic lobes were embedded in 4% agarose with low melting temperature (GTG-NuSieve Agarose, Lonza) before being sectioned into 200-400□m slices using a vibratome (1000 Plus sectioning system, Leica). Thymic slices were overlaid onto 0.4□m transwell inserts in 6 well tissue culture plates containing 1ml of complete RPMI medium (containing 10% FBS, penicillin streptomycin, and 2-mercaptoethanol, cRPMI) under the transwell, and cultured at 37°C. 10□1 of cRPMI containing 0.5-2×10^6^ thymocytes was overlaid onto each slice, and thymocytes were allowed to migrate into the slice for 2 hours following the removal of excess thymocytes by gentle washing of the slice with PBS. Thymocytes actively migrate into the slices and localize properly within the tissue^28, 56^. Overlaid thymocytes were distinguished from slice-endogenous thymocytes using CD45.1/CD45.2 congenic markers, or in some cases, by staining with cell proliferation dyes. To introduce cognate antigen, 10□1 of 1□M SIINFEKL (AnaSpec) or SIIQFEKL (Q4, AnaSpec) in PBS was overlaid onto each slice for 30 minutes before excess was removed by pipetting. For IL-15 treatment, slices were overlaid with 50ng recombinant IL-15/IL-15Rαcomplex (Thermo Fisher) in 10□1 of cRPMI following peptide treatment. To make IL-2 complexes, 1.5□g of recombinant IL-2 (Peprotech) and 7.5□g of functional grade anti-IL-2 antibody (JES6-1, Thermo Fisher) were incubated together at 37°C for 30 minutes. Thymic slices were treated with 75ng IL-2 (in complex) in 10□1 PBS following peptide treatment.

### Bone marrow-derived dendritic cell cultures and *in vitro* activation

Bone marrow was collected from the femurs and tibias of mice and resuspended in ammonium chloride–potassium bicarbonate buffer for lysis of red blood cells. Bone marrow cells were then plated at 10^6^ cells/ml in cRPMI containing 20ng/ml granulocyte-macrophage colony-stimulating factor (GM-CSF, Peprotech), and cultured for 7 days. On day 6, media was replaced with fresh media containing GM-CSF. Semi-adherent cells were collected at day 7 and loaded with SIINFEKL in cRPMI containing 1□M peptide at 10^7^ cells/ml for 30 minutes at 37°C. BMDCs were then plated at 2×10^5^ cells per well in 96 well U-bottom tissue culture plates and 3×10^5^ thymocytes stained with 0.5□M eFluor670 (Thermo Fisher Scientific) were added per well. BMDCs and thymocytes were cultured together for 48 hours prior to collection for flow cytometric analysis.

### Adoptive transfers and isolation of intraepithelial lymphocytes

Thymic slices were collected at the indicated timepoints, dissociated, and filtered to remove agarose. 3×10^6^ total cells were then injected intravenously into Rag2^-/-^ mice 4-7 weeks of age, and intraepithelial lymphocytes were collected from the small intestine at 7 days post-transfer. The small intestine was harvested and cleaned of fat and connective tissue. Fecal contents were flushed out with PBS, and the tissue was cut into small pieces that were placed in 1mM DTT in Hank’s Balanced Salt Solution, and shaken at 37°C for 1 hour total, with collection of cells and replacement of new 1mM DTT solution every 20 minutes. Samples were resuspended in 20% Percoll and overlaid onto 40% Percoll over 70% Percoll and spun. The 40/70 interface was collected and stained for flow cytometric analysis.

### Flow cytometry

Single cell suspensions were resuspended in 24G2 supernatant containing the following antibodies: CD4 (GK1.5), CD8α̣(53-6.7), CD8β̣(H35-17.2), CD69 (H1.2F3), CD122 (Tm-b1), α4β7 integrin (DATK32), PD-1 (29F.1A12), CD3 (145-2c11), TCRβ̣(H57-597), CD45.1 (A20), CD45.2 (104), CD28 (37.51), on ice for 10 minutes. All antibodies were from Tonbo Biosciences, Thermo Fisher Scientific, or Biolegend. Cells were then washed in PBS and stained in Ghost Dye Violet 510 (Tonbo Biosciences) for 10 minutes on ice. To obtain single cell suspensions from thymic slices, slices were dissociated into FACS buffer (0.5% BSA in PBS) and filtered for removal of agarose before staining. For intracellular staining of IFNγ, cells were cultured with Protein Transport Inhibitor Cocktail for 4 hours at 37°C prior to staining. Following staining with cell surface antibodies and live/dead staining, cells were fixed using Cytofix/Cytoperm Kit (BD Pharmingen), then stained for IFNγ (XMG1.2). For intracellular staining of T-bet, cells were stained with cell surface antibodies and live/dead dye, then fixed and permeabilized using the FoxP3 Transcription Factor Staining Buffer Set (Thermo Fisher Scientific) according to the manufacturer’s protocol prior to staining for T-bet (644814). All data was collected on a LSR II or BD Fortessa analyzer (BD Biosciences), and data were analyzed using FlowJo software (TreeStar).

### Microscopy

For labeling, 10^7^ thymocytes were incubated with 2 µM Indo-1 LR (Teflabs) at 3X10^6^ cells/ml for 90 min in cRPMI at 37°C. For overlaying on thymic slices, the cell suspensions were adjusted to 5×10^5^ cells/10 µl and 10µl were gently overlaid onto thymic slices in transwells as described above. The cells were left to migrate into the slice for 2 hours at 37°C/5% CO2, followed by gentle washing with PBS to remove the cells that failed to enter the slice. Thymic slices were prepared from RIPmOVA transgenic or non-transgenice mice. For some experiments RIPmOVA mice were crossed to AIRE-GFP^54^ or CD11cYFP^53^ reporter mice to allow for visualizing mTEC or dendritic cells, respectively. Thymic tissue slices were glued on coverglasses and medullary regions, identified by density of CD11cYFP or AIRE-GFP reporter+ cells, were imaged by two-photon laser scanning microscopy with a Zeiss 7 MP (Zeiss), while being perfused with warmed (37°C), oxygenated phenol-free DMEM medium (GIBCO) at a rate 1 ml/min. Mode-locked Ti:sapphire laser Mai-Tai (Spectra-Physics) or Chameleon (Coherent) was tuned to 720nm for Indo-1 LR excitation, or to 920nm for CFP or GFP excitation, with appropriate filter sets. Imaging volumes of various sizes were scanned every 30 sec for 20-60 min.

### Image analysis

Imaris 7.3 (Bitplane) was used to determine cell positions over time and tracking. The x, y and z coordinates as well the mean fluorescence intensities of the tracking spots for Ca^2+^-bound Indo-1 LR, Ca^2+^-free Indo-1 LR were exported. Motility parameters were calculated in MATLAB (Mathworks) with a custom code that is available upon request. Ca^2+^-ratio was calculated as a surrogate for Ca^2+^ intracellular concentration by dividing the mean fluorescence intensity of Ca-^2+^-bound Indo-1 LR by the Ca^2+^-free Indo-1 LR. All the values were normalized so that the average Ca^2+^-ratio of migrating thymocytes in the no Ova control sample was zero (corrected Ca^2+^-ratio).

### Statistics

Prism software (GraphPad) was used for all statistical analysis. P values of <0.05 were considered significant.

## ACKNOWLEDGEMENTS

We thank members of the Robey lab for helpful discussion and technical advice, and H. Melichar for critical reading of the manuscript. We also thank S.W. Chan and O. Guevarra for technical assistance, P. Herzmark for assistance with two-photon imaging, and M. Anderson (UCSF) for providing AIRE-GFP reporter mice. This work was funded by NIH RO1AI064227 (E.A.R.). N.S.K. and B.MW. were supported by NIH T32AI100829, and N.S.K. was supported by a University of California Cancer Research Coordinating Committee Fellowship.

## AUTHOR CONTRIBUTIONS

N.S.K. designed and performed experiments, analyzed data and wrote the manuscript.

A.H. designed and performed experiments and analyzed data.

J.Y. performed experiments and analyzed data.

B.M.W. designed and performed experiments.

E.A.R. designed experiments, analyzed data and wrote the manuscript.

---

**Supplementary Movie 1. Migration of OT1 thymocytes in confinement zones: example 1.**

OT-I thymocytes were labeled with the ratiometric fluorescent calcium indicator dye IndoPE and overlaid onto thymic slices prepared from RIPmOVA mice crossed to CD11c-YFP reporter mice, and imaged by 2 photon microscopy after 3 hours of culture. Time lapse image show signal from IndoPE, and rotation views show the thymocyte tracks superimposed on the YFP signal (white) collected at the end of the time-lapse sequence. Imaging dimensions are 87 × 87 × 104 microns × 30 minutes.

**Supplementary Movie 2. Migration of OT1 thymocytes in confinement zones: example 2.**

As in Movie 1, except that thymic slices were prepared from RIPmOVA mice crossed to AIRE-GFP reporter mice. Time lapse image show signal from IndoPE, and rotation views show the thymocyte tracks superimposed on the GFP signal (white) collected at the end of the time-lapse sequence. Imaging dimensions are 149 × 149 × 84 microns × 30 minutes.

