## Supplementary Figures for "Factors that influence the thymic selection of CD8αα intraepithelial lymphocytes"

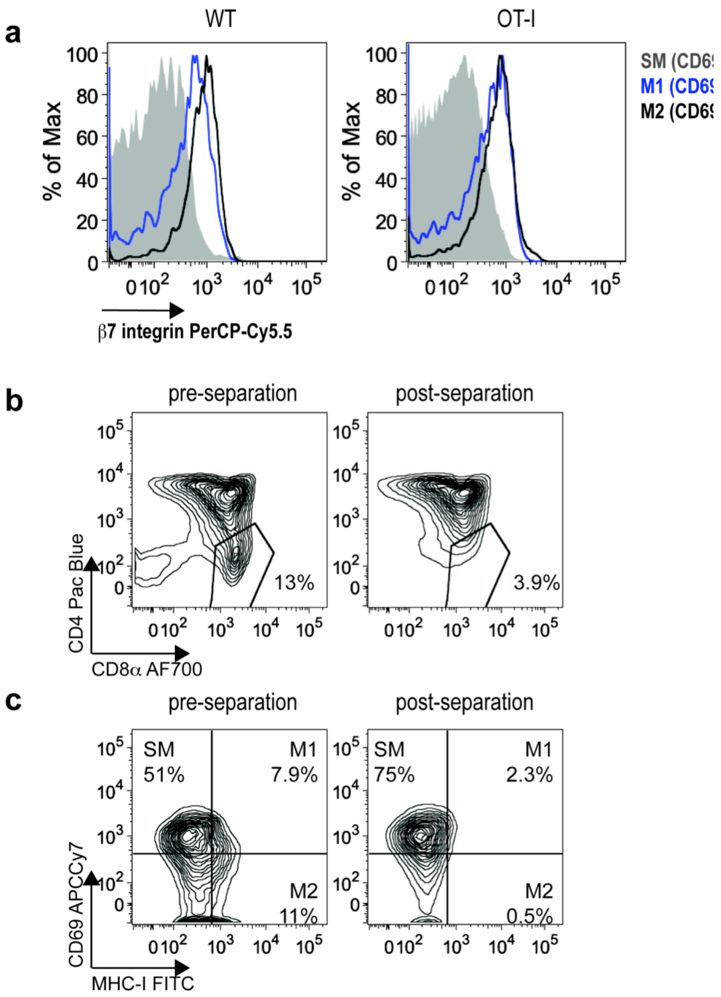


**Supplementary Figure S1. β7 integrin depletion removes the most mature CD8 single positive thymocytes**

(a) Expression of β7 integrin on TCRβ+CCR7+ WT or OT-I thymocyte subsets of increasing maturation: semi-mature (SM, CD69+MHC I+, gray), M1 (CD69+MHC I-, blue), and M2 (CD69-MHC I-, black){Hogquist:2015fc}. (b) Representative flow cytometry plots displaying the proportion of CD8 single positive (CD8 SP) OT-I thymocytes before (left) or after (right) β7 integrin depletion. (c) Representative flow cytometry plots displaying the proportions of SM, M1, and M2 OT-I thymocytes before (left) or after (right) β7 integrin depletion. Data are representative of 2 independent experiments.

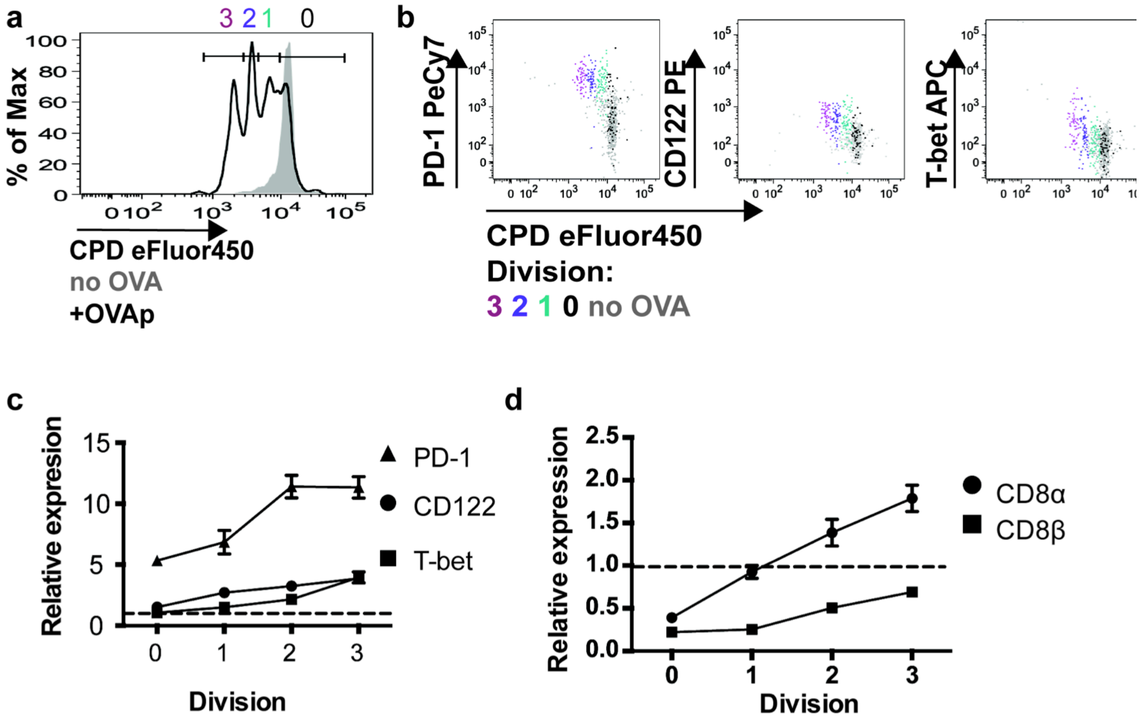


**Supplementary Figure S2. Phenotypic changes in thymocytes as they proliferate in response to cognate antigen**

OT-I thymocytes depleted of mature CD8 single positives were stained with CPD450, overlaid onto thymic slices to which OVAp was added, and harvested 48 hours later for flow cytometric analysis. (a) Dilution of CPD450 in OT-I thymocytes, with gates to determine the number of cell divisions undergone shown. (b) PD-1, CD122, or T-bet expression relative to cell proliferation, displayed as representative flow cytometry plots. (c,d) Expression of PD-1, CD122, and T-bet (c), or CD8α and CD8β (d), in cells that have not divided, or divided 1, 2, or 3 times (as determined by dilution of CPD450, gates shown in a), displayed as Mean Fluorescence Intensity (MFI) relative to MFI in no OVA control slices, with mean and SEM shown. Data are compiled from (c,d) or representative of (a,b) 3 independent experiments, with n=14 individual slices per condition.

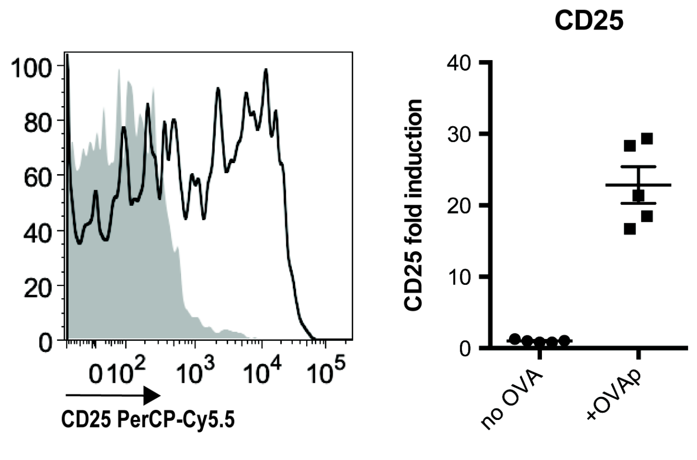


**Supplementary Figure S3. Thymocytes upregulate CD25 upon antigen encounter**

OT-I thymocytes were overlaid onto thymic slices to which OVAp was added, and slices were harvested for flow cytometric analysis 16 hours later. Expression of CD25 presented as representative flow cytometry plot of gated OT-I thymocytes (left), or fold induction displayed as MFI of CD25 relative to levels in no OVA control slices (right). Data are representative of (left), or pooled from (right) n=5 individual thymic slices, with mean and SEM, where each dot represents an individual slice. ****$p<0.0001$ (unpaired t test)

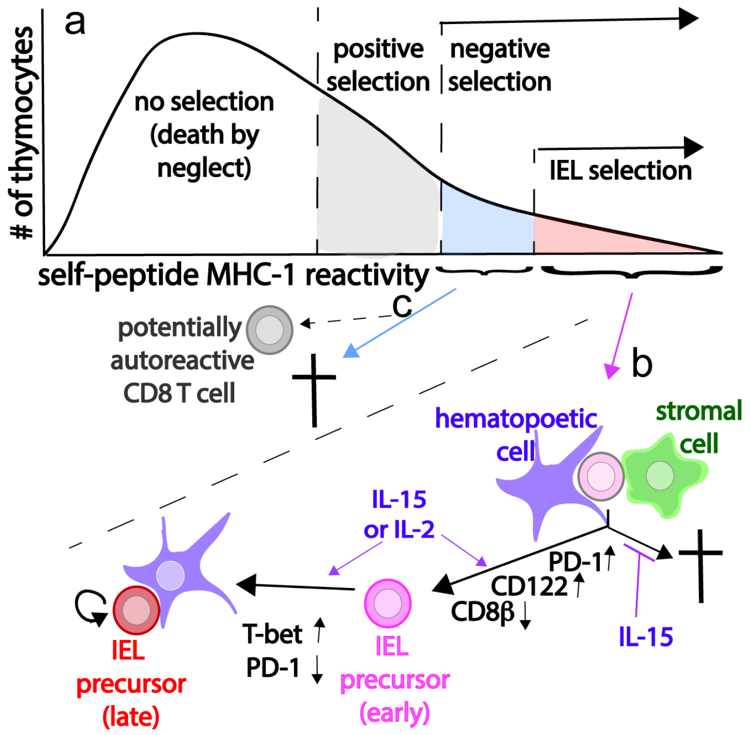


**Supplementary Figure S4. Model for the development of CD8αα IEL precursors.** (a) Hypothetical distribution of levels of thymocyte self-reactivity based on the ability of the TCR to engage self-peptide MHC class I complexes. Low to moderate self-reactivity leads to positive selection (grey), moderately high reactivity can trigger negative selection (light blue), and very high self-reactivity can trigger CD8αα IEL development (pink). (b) Thymocytes that encounter very high avidity self-peptides presented by hematopoietic or non-hematopoietic APC initiate CD8αα IEL development, including transient PD-1 upregulation, CD122 upregulation, and CD8β down-regulation. IL-15 protects cells from negative selection, while both IL-15 and IL-2 can promote IEL development (purple arrows). Additional interactions with hematopoietic APC can lead to proliferation and T-bet induction. (c) While most class I MHC specific thymocytes with moderately high self-reactivity undergo negative selection, some may escape, posing a hazard for autoimmunity.
